# Adaptive divergence in transcriptome response to heat and acclimation in *Arabidopsis thaliana* plants from contrasting climates

**DOI:** 10.1101/044446

**Authors:** Nana Zhang, Elizabeth Vierling, Stephen J. Tonsor

## Abstract

Phenotypic variation in stress response has been widely observed within species. This variation is an adaptive response to local climates and is controlled by gene sequence variation and especially by variation in expression at the transcriptome level. Plants from contrasting climates are thus expected to have different patterns in gene expression. Acclimation, a pre-exposure to sub-lethal temperature before exposing to extreme high temperature, is an important adaptive mechanism of plant survival. We are interested to evaluate the gene expression difference to heat stress for plants from contrasting climates and the role of acclimation in altering their gene expression pattern. Natural *Arabidopsis thaliana* plants from low elevation mediterranean and high elevation montane climates were exposed to two heat treatments at the bolting stage: a) 45°C: a direct exposure to 45°C heat; b) 38/45°C: an exposure to 45°C heat after a 38°C acclimation treatment. Variation in overall gene expression patterns was investigated. We also explored gene expression patterns for Hsp/Hsf pathway and reactive oxygen species (ROS) pathway. In both heat treatments, high elevation plants had more differentially expressed (DE) genes than low elevation plants. In 45°C, only Hsp/Hsf pathway was activated in low elevation plants; both Hsp/Hsf and ROS pathways were activated in high elevation plants. Small Hsps had the highest magnitude of change in low elevation plants while Hsp70 and Hsp90 showed the largest magnitude of fold in high elevation plants. In 38/45°C, Hsp/Hsf and ROS pathways were activated in both low and high elevation plants. Low elevation plants showed up-regulation in all Hsps, especially small Hsps; high elevation plants showed down-regulation in all Hsps. Low elevation and high elevation also adopted different genes in the ROS pathway. We also observed genes that shifted expression in both low and high elevation plants but with opposite directions of change. This study indicates that low and high elevation plants have evolved adaptive divergence in heat stress response. The contrasting patterns of temperature variation in low and high elevation sites appears to have played a strong role in the evolution of divergent patterns to high temperature stress, both pre-acclimation and direct exposure gene expression responses.

**Molecular Ecology:** The Plant Journal IF: 6.8 (TPJ welcomes functional genomics manuscripts when a scientific question, rather than the technology used, has driven the research)

## Introduction

Abiotic stresses are major driving forces in evolutionary diversification (Badyaev 2005; Hoffmann & Hercus 2000; Hoffmann & Parsons 1991). Diversification in adaptation to environments with contrasting patterns of stresses is important in shaping ecological structure in nature (Keller & Seehausen 2012).

Plants have evolved various abiotic stress response mechanisms at morphological, physiological and biochemical levels, with diversity in responses evidenced both within and between species (Berry & Bjorkman 1980; Yeh *et al.* 2012; Zhang *et al.* 2015a). Local adaptation to stressful environments has been extensively explored, such as adaptation to drought stress (Bowman *et al.* 2013; McKay *et al.* 2003; Shinozaki & Yamaguchi-Shinozaki 2007; Zhu 2002), salt stress (Zhao *et al.* 2012; Zhu 2002), cold stress (Beales 2004; Sakai & Larcher 1987; Shinozaki *et al.* 2003), and heat stress (Kotak *et al.* 2007; Rizhsky *et al.* 2002; Rizhsky *et al.* 2004b). However, few studies have explored the transcriptional variation underlying variation in phenotypic stress responses. Identification of changes in gene expression involved in diversification of abiotic stress responses is an important step in understanding the evolutionary response to stress-mediated natural selection. Furthermore, understanding evolved variation in gene expression response to stresses at a population level can provide insight on the cause of the capacity/limit of an organisms’ ability to adapt to local climate and the mechanisms by which differential adaptation has been effected. We are particularly interested in the adaptive response of heat stress due to its increasing importance in global climate change events.

Heat stress response involves large scale gene reprograming at the level of the transcriptome in the context of complex regulatory networks (Dittami *et al.* 2009; Liu *et al.* 2013a). The multiple genes discovered by RNA-seq analysis among animal and plant species have suggested a complicated structure to the response to heat stress (Kotak *et al.* 2007). Two main pathways are activated during exposure to heat (Ahuja *et al.* 2010; Kotak *et al.* 2007; Qu *et al.* 2013). Heat stress first activates the up-regulation of heat shock transcription factors (Hsfs) and heat shock proteins (Hsps) (Baniwal *et al.* 2004; Wang *et al.* 2004). The highly conserved Hsps are the most extensively studied heat stress related genes. Oxidative stress, as a secondary stress, is also activated during heat stress (Qu *et al.* 2013). Reactive oxygen species (ROS) pathway, the expression of transcription factors in Zat and WRKY family, MBF1c and Rboh, is thus activated (Ciftci-Yilmaz *et al.* 2007; Rizhsky *et al.* 2004a; Suzuki *et al.* 2008; Suzuki & Mittler 2006). The response involves several critical biological processes, such as antioxidant system neutralization of free radicals, protein synthesis and degradation, plant hormone production (e.g. salicylic acid) (Liu *et al.* 2013a; Liu *et al.* 2013b; Narum & Campbell 2015; Qu *et al.* 2013).

The adaptive responses to climate variables are highly dependent on the geographic origin of the populations and their genetic background (Schimper 1902). Gene expression response to the application of the plant hormone salicylate varied in *Arabidopsis thaliana* populations from diverse climate origins (Leeuwen *et al.* 2007). Our study region, the northeastern Iberian Peninsula, provides an ideal location for studying the general patterns of response to climate in plants. Populations collected across an elevation gradient provide a platform to examine adaptation to diverse climates (Clausen & Hiesey 1958; Schimper 1902). Two major climate variables, annual temperature and precipitation, follow closely with elevation along a gradient from the Mediterranean coast into the Pyrennee mountains (Wolfe & Tonsor 2014). Importantly, native *Arabidopsis thaliana* populations in this region show morphological and physiological divergence, such as life cycle timing (Wolfe & Tonsor 2014), seed dormancy and germination traits (Montesinos-Navarro et al. 2012) and one key plant hormone, salicylic acid (Zhang *et al.* 2014).These native *Arabidopsis* populations also show divergent response to various abiotic and biotic stresses. For example, sixteen populations showed differential Hsp101 expression when exposed to a 42°C compared to a 45°C heat treatment (Zhang *et al.* 2015a), while salicylic acid differed among four tested populations when exposed to a 44°C heat treatment for 3hrs(Zhang *et al.* 2015b). These populations also show differential expression when exposed to cold stress and pathogen infection (Zhang *et al.* 2014). Recently, adult plants under heat stress showed contrasting avoidance and tolerance strategies when comparing plants from contrasting climates (Zhang *et al.* under review). These diverse and contrasting strategies for low vs. high elevation plants suggest that adaptive and fine-tuned heat stress mechanisms are involved.

Acclimation, a process resulting from a pre-exposure to sub-lethal high temperature before exposing plants to the extreme high temperature, is an important adaptive mechanism of plant survival when in a high temperature environment (Alscher & Cumming 1990; Badger *et al.* 1982; Whitehead 2012). A previous microarray experiment from Larkindale and Vierling (2008) showed that two heat treatment regimes, one with a moderately high temperature acclimation followed by high temperature, the other a direct exposure to high temperature, have very different core genome responses (Larkindale & Vierling 2008). Both the number and abundance of transcripts up-regulated and down-regulated under heat stress (compared to the control condition) differ between the two heat treatment regimes. In addition, among the multiple genes that are involved in acclimation to high temperature, there appears to be more than one strategy that achieves similar protective effects (Larkindale & Vierling 2008). Thus in our study, we looked the transcriptome response to heat stress with or without an acclimation treatment. Identifying the specific gene set in each heat stress regime and elucidating the complete mechanisms of heat stress response will contribute to fine-scale control for future breeding programs as well as for predicting the response to future climate change.

RNA-Seq has become a powerful and revolutionary tool to investigate the divergent responses to various thermal climates within species when they are exposed to the same heat stress (Wang *et al.* 2009). In this study, our goals were to identify whether/how plants from contrasting climates showed different gene expression patterns in response to heat and whether/how an acclimation treatment altered the gene expression patterns. To do this, we exposed low and high elevation *Arabidopsis thaliana* plants to two 45°C heat treatments: one without and one with a 38°C acclimation. We firstly compared the constitutive gene expression level between the low and high elevation plants in the control. We then identified elevation specific significantly differentially expressed (DE) genes by contrasting low and high elevation plants within each treatment (within heat treatment, across elevation groups). We identified the elevation specific DE genes for both treatments. Next we identified acclimation specific DE genes by comparing the two heat treatments for each elevation group (i.e. within elevation group, across heat treatment). We specifically looked into the gene expression level of currently known heat stress related DE genes, including heat shock proteins (Hsps), heat shock transcription factors (Hsfs) and many others, in our elevation specific and acclimation specific DE genes. We also investigated the functions of the genes were DE for both low and high elevation plants but with opposite directions of changes for the plants from the two climate regions. We did this for each heat treatment. Our study shed light on evolutionary adaptation to local climates, especially past high temperature events, at the transcriptome level.

## Materials and methods

### *Arabidopsis thaliana* materials and treatments

Plants from eight populations, four from low and four from high elevation, were chosen as representative plants. Since *Arabidopsis thaliana* is highly selfing and highly genetically homogenous within populations (Tang *et al.* 2007), we selected four plants, one genotype per population, to represent the plants in each elevation region. To test for differential responses to heat and the role of acclimation, we designed two heat treatments that we compared to a control group. 24 plants total, consisting of three replicates of each of the eight unique plants, were blindly divided into the three treatment groups. All plants were germinated following a five-day 4°C stratification in the dark and maintained at 22°C for three weeks (16 hrs light/8 hrs dark) in Conviron PGW36 controlled environment growth chamber (http://www.conviron.com) at the University of Pittsburgh. After three weeks of growth, seedlings then experienced a four-week vernalization treatment at 5°C to synchronize flowering time.

Since these plants are most likely to experience heat stress at the bolting stage in nature, heat treatments were imposed at standard stage 6.0-6.1 (Boyes *et al.* 2001). Following vernalization, plants were observed daily and those at the stage 6.0-6.1 were selected for heat treatment in a separate PGW36 chamber. The heat treatments were: a) 45°C: a 45°C treatment for 3hrs; and b) 38/45°C: a 38°C acclimation for 3hrs followed 2hrs later with a 3hr 45°C treatment. The control group was maintained at a constant 22°C throughout the experiment. Each treatment group included all eight plants. After placement of plants in the heat treatment chamber, the temperature increased over a 15 minute period from a starting temperature of 22°C, as in Larkindale and Vierling, 2008.

### RNA extraction, cDNA library construction and RNA sequencing

Leaf samples for both heat treatment and control plants were collected immediately after the heat treatment, stored in liquid nitrogen and transferred to a −80°C freezer. After leaf samples were freeze-dried, RNA was extracted and purified using Qiagen RNeasy Plant Mini Kit (Qiagen) using the kit’s instruction manual recommended protocol. The quality and quantity of the RNA samples were measured using Qubit Fluorometric quantitation (Thermo Fisher Scientific). Total RNA samples were then adjusted to a concentration of 100ng/ul in 25ul nuclease free water (2.5 ug total) for cDNA library construction. Before cDNA library construction, all RNA samples were evaluated via Bioanalyzer for further RNA quality assessment (Genomics Research Core, Health Science Core Research Facilities, University of Pittsburgh). RNA samples were re-extracted and re-purified if they did not pass the quality control.

Next the poly-A RNAs were converted into ds-cDNA and fragmented into 100bp fragments. cDNAs were then ligated with adaptors and amplified with PCR. The cDNA libraries were constructed using the Truseq RNA Sample Prep kit (Illumina) in the DNA Core, University of Missouri.

Eight cDNA libraries were combined per pool, three pools total. Each pool was sequenced in a single lane of a 1x100bp single-end Illumina HiSeq 2000 run, 3 lanes total. The sequencing was done in November 2014 at the University of Missouri DNA Core. The aligned RNA sequences have been uploaded to the NCBI sequences database (accession number xxxx).

### RNA sequence mapping, differential expression and functional categorization

Raw read data were first checked with FastQC software for quality control. Because the per base QC content was high for the first 15bp, the sequences were processed with FastX Trimmer to trim the first 15bp and last 15bp for each 100bp sequence for high quality sequence alignment.

The reads were then mapped to *Arabidopsis thaliana* Col-0 (Tair-10) transcriptome with Tophat2. Two mismatches were allowed in each segment alignment for reads mapped independently. We obtained an average of 28M raw reads per sample. An average of 23.7M reads, 85.8% of the raw reads, were mapped to the reference genome (Table S1).

To understand the constitutive gene expression between low and high elevation plants, we firstly identified the up-and down-regulated significantly differentially expressed (DE) genes by contrasting high elevation plants with low elevation plants in the control. The comparison and the functional annotation was performed using CuffDiff2 with p = 0.01 (Trapnell *et al.* 2012).

Next, to categorize the DE genes, we adopted two approaches. For both approaches, we firstly calculated the difference in gene expression between each heat treatment and the control, separately for low or high elevation plants. The difference indicates differential gene expression (DE) response to the heat treatments. Our first approach was to contrast the gene expression response of low and high elevation plants to each heat treatment. The second approach was to compare the gene expression in 45°C and 38/45°C to investigate the role of 38°C acclimation. We contrasted the gene expression of 45°C and 38/45°C within elevation groups, comparing for low and high elevation plants respectively. The detection of DE genes, their fold change, and normalized fold change (FPKM-Fragments Per Kilobase of Exon per Million Fragments Mapped), as well as their functional annotations, were conducted using Cuffdiff2 with p value = 0.01 (Trapnell *et al.* 2012). For all comparisons, we firstly contrasted all the DE genes regardless of the direction of change. We then compared up-regulated and down-regulated DE genes separately. By doing the above comparison, we also uncovered DE genes that exhibited opposite direction of change between elevation groups. The DE shared across elevation groups and the DE genes that were unique to an elevation group were compared and visualized using BioVenn, a web application (http://www.cmbi.ru.nl/cdd/biovenn/index.php). The functional categorization of the DE genes was performed in TAIR (https://www.arabidopsis.org/tools/bulk/go/index.jsp).

We furthered explored the elevation specific and acclimation specific DE genes from the above comparisons. We investigated the currently known stress-related genes in the Hsp/Hsf pathway and the ROS pathway from a literature review list (Table S2). We used these known heat stress related genes and their magnitude of change in heat to represent the elevation or acclimation specific response in our study. In our approach of comparing DE genes regardless of direction with separate up-and down-regulated DE gene, we also uncovered 51 shared DE genes DE between high and low elevation plants, but with directions of change (35 DE genes in 45°C, 19 DE genes in 38/45°C, with three shared DE genes; Table 4). Their function and possible biological processes involved were also double checked in NCBI database (http://www.ncbi.nlm.nih.gov).

## Results

### Constitutive gene expression difference in low vs. high elevation plants

When expression levels in high elevation plants were compared to low elevation plants in the control, 1291 DE genes were found. Of these, 826 were up-regulated and 465 were down-regulated in the high elevation plants relative to low elevation plants. We found eight Hsps, including Hsp60, Hsp70 and Hsp90 family, that showed up-regulation in high elevation plants (Table 1). Zat7 and Zat10, responding to various stresses, were also up-regulated in high elevation plants. ABI2, involved in abscisic acid (ABA) signaling, NDH1, providing protection against photo-oxidation, and FtsH11, associated with reduced photosynthetic capacity in heat stress, were down-regulated in high elevation plants. Hsps respond via Hsp/Hsf pathway, Zat genes respond to ROS pathways. The up-regulation in Hsps and Zat indicates that high elevation plants were constitutively more resistant to heat stress. Down-regulation in NDH1 and FtsH11 indicates a negative control in photosynthesis in high elevation plants.

**Table 1.** Constitutive expression difference in currently known heat stress related genes comparing high to low elevation plants

| Gene ID | Gene Name | FPKM |
| --- | --- | --- |
| AT5G56030 | Hsp90-2 | 292.6 |
| AT3G12580 | Hsp70 | 110.86 |
| AT3G23990 | Hsp60-3B | 78.1 |
| AT3G07770 | Hsp89.1 | 36.33 |
| AT2G33210 | Hsp60-2 | 23.65 |
| AT4G21870 | Hsp class V 15.4 | 127.13 |
| AT4G30350 | Hsp | 58.65 |
| AT5G51440 | Hsp23.5 | 4.92 |
| AT3G46090 | Zat7 | 30.57 |
| AT1G27730 | Zat10 | 671.6 |
| AT1G21910 | DREB26 | 17.72 |
| AT5G57050 | ABI2 | -25.22 |
| AT1G15980 | NDH1 | -310.35 |
| AT5G53170 | FtsH11 | -78.47 |

Note: FPKM, short for Fragments Per Kilobase Of Exon Per Million Fragments, is the normalized fold changes in gene expression when comparing high elevation plants to low elevation plants. Positive value means up-regulation, and negative value means down-regulation.

In plants, MADS-box genes play major roles in controlling development and determining flowering time. The MADS-box gene FLOWERING LOCUS C (FLC) and SOC1 (AGL20, SUPPRESSOR OF OVEREXPRESSION OF CONSTANS1) are necessary for the correct flowering timing. In high elevation relative to low elevation plants, FLC gene was up-regulated and the SOC1 was down-regulated. Another MADS-box gene AGL3 was also down-regulated. The constitutive gene expression difference in MADS box genes indicates inherent variation in flowering time between low and high elevation plants.

### Elevation-specific DE genes in response to heat

When exposed directly to a 45°C heat, 1516 and 2104 DE genes were found for low and high elevation plants, respectively (Fig.1a). Similarly, when exposed to the 38/45°C heat, 1489 and 1972 DE genes were found for low and for high elevation plants, respectively (Fig.1b). High elevation plants showed 39% and 32% more DE genes than low elevation plants in 45°C and 38/45°C heat, respectively. High elevation plants in the 45°C heat showed the most DE genes. We further compared the magnitude of fold change in low and high elevation plants in the two heat treatments (Fig.2). High elevation plants in the 45°C heat also showed the largest average magnitude of fold change among the four bars. The average magnitude of fold change was similarly low.

**Figure 1.**
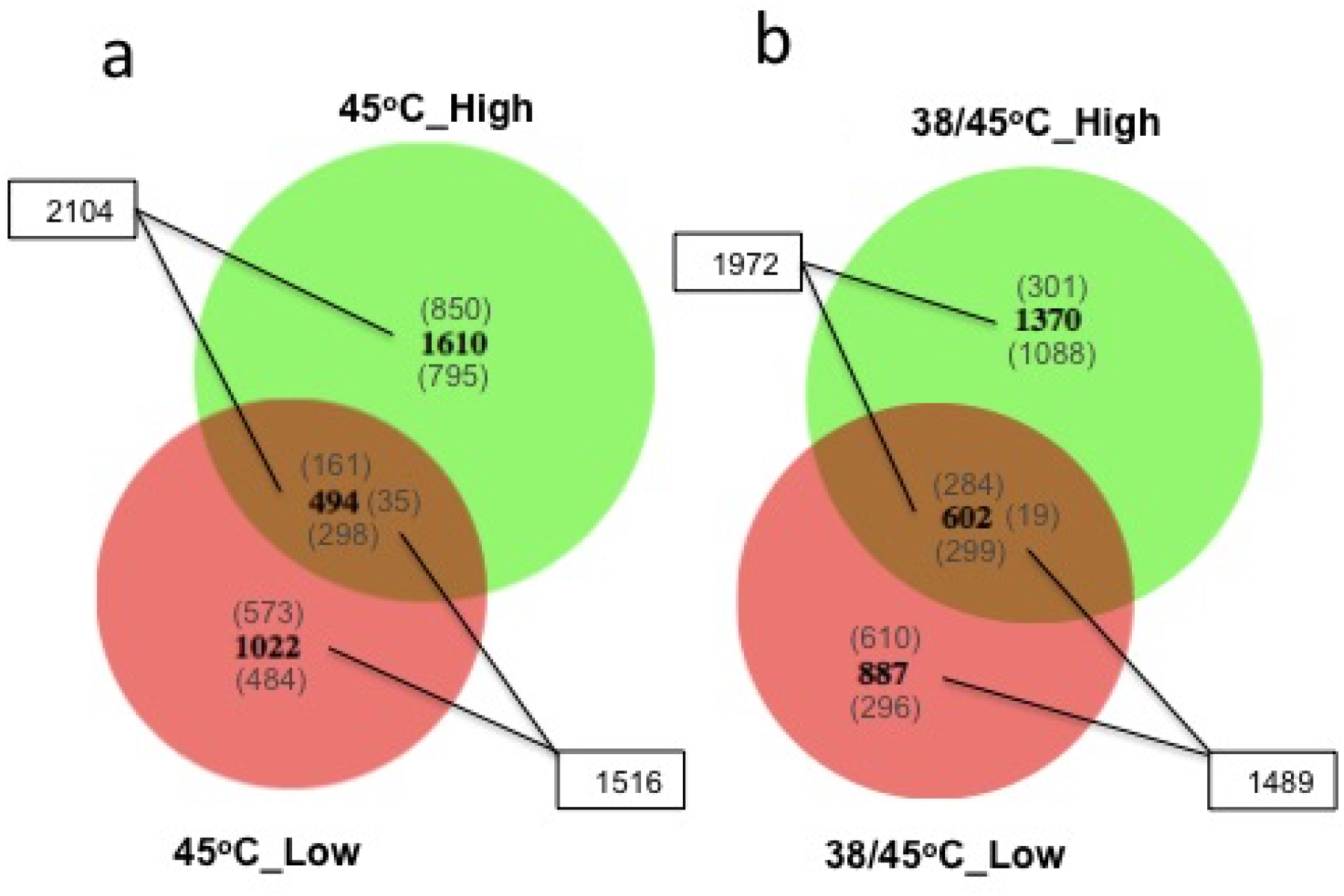
Venn diagrams comparing the number of DE (significantly differentially expressed) genes for the indicated treatment and elevation groups. In all cases DE genes are those differing significantly in expression compared to the same genes expressed in the corresponding 22°C control group. Bold numbers indicate total number of genes showing changed expression. Numbers in parentheses above the bold number indicate up-regulated genes and numbers below the bold indicate those that are down-regulated. The area of overlap of the two circles indicates the proportion of DE that are shared between the treatment and elevation groups. In the area of overlap, numbers in parentheses to the right of the bold numbers indicate DE genes that were shared but with directions of change between the compared groups, e.g. up-regulated in one group but down-regulated in the other group.

**Figure 2.**
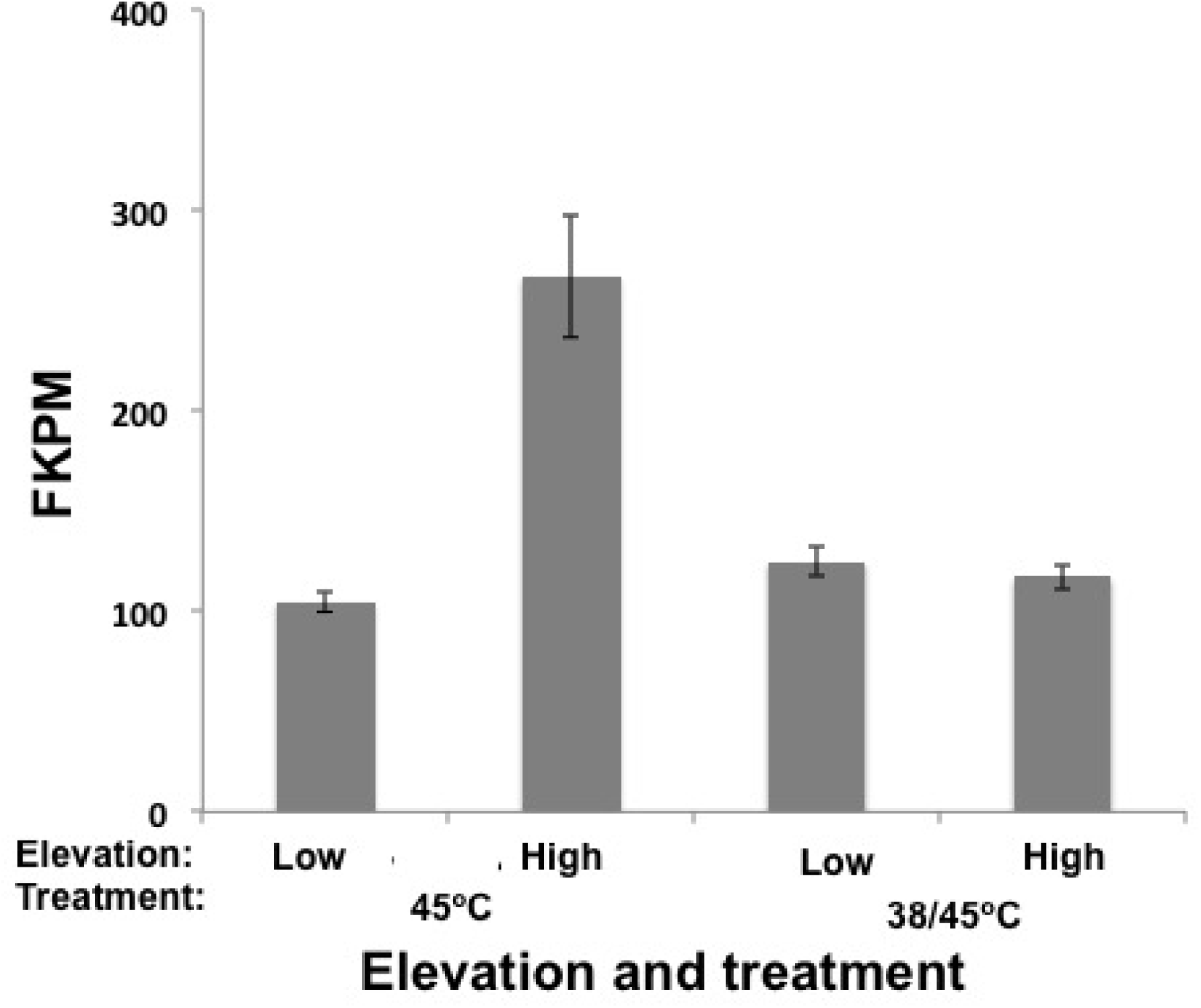
The difference in magnitude of the normalized fold change, FKPM, of the DE genes. The data showed are means of FKPM value for each treatment elevation pair. Error bars are standard errors.

When the DE genes in the 45°C heat were compared between the two elevation groups, 494 shared DE genes were identified. Among these 494 shared DE genes, 161 were up-regulated, 298 were down-regulated, and 35 showed directions of change. For the elevation specific DE genes, high elevation plants had 58% more uniquely DE genes than low elevation plants (1610 vs. 1022, Fig.1a). Similarly, we found 602 shared DE genes between low vs. high elevation plants after the 38/45°C heat. 284 of these were up-regulated, 299 were down-regulated, and 19 showed opposite directions of change in the two elevation groups. High elevation plants have 54% more unique DE genes than low elevation plants (1370 vs. 887) (Fig.1c). We further compared the numbers of up-regulated and down-regulated DE genes between the two elevation groups. In 45°C, both low and high elevation plants showed more elevation-specific up-regulated (573 and 850) than down-regulated DE genes (484 and 795). However, in 38/45°C, there were more high elevation specific down-regulated DE genes (1088) than up-regulated DE genes (301), while more low elevation-specific up-regulated (610) than down-regulated DE genes (296) (Fig.1).

To further explore the elevation-specific DE genes, we investigated the expression level of each gene in Table S1 from the above-mentioned elevation-specific DE genes (Fig.1), for each heat treatment (Table 2). Genes in Table S1 include genes involved in Hsp/Hsf pathway and ROS pathway. In 45°C, we found 10 low elevation-specific and 22 high elevation-specific DE heat stress related genes. In low elevation plants, several small Hsps, such as Hsp20, Hsp21, Hsp22, had the largest up-regulation; while in high elevation plants, the Hsp70 and Hsp 90 sub-families had the largest up-regulation. Low elevation plants also differentially expressed Hsp60, Hsp70, and Hsp90 sub-family genes, but with different DE genes within the sub-families compared to high elevation plants and with significantly lower magnitude of fold change. Hsp101, the only Hsp known to be necessary for acquired thermotolerance (Hong & Vierling 2001), was uniquely expressed in low elevation plants only. Only one Hsf showed down-regulation in low elevation plants but five Hsfs showed up-regulation in high elevation plants, High elevation plants also showed DE in eight ROS related genes. DREB2 and DREB2B were up-regulated. Up-regulation of DREB genes activates the expression of stress-related genes. RBohD and RBohF were ROS signal amplifiers and were down-regulated. DGD1 and DGD2, whose expression were associated with reduction in photosynthetic capacity, were up-regulated. To summarize, in low elevation plants, only the Hsp/Hsf pathway was activated and small Hsps had the highest magnitude of change; in high elevation plants, both Hsp/Hsf and ROS pathways were activated, with Hsp70 and Hsp90 showing the largest magnitude of fold change.

**Table 2.**
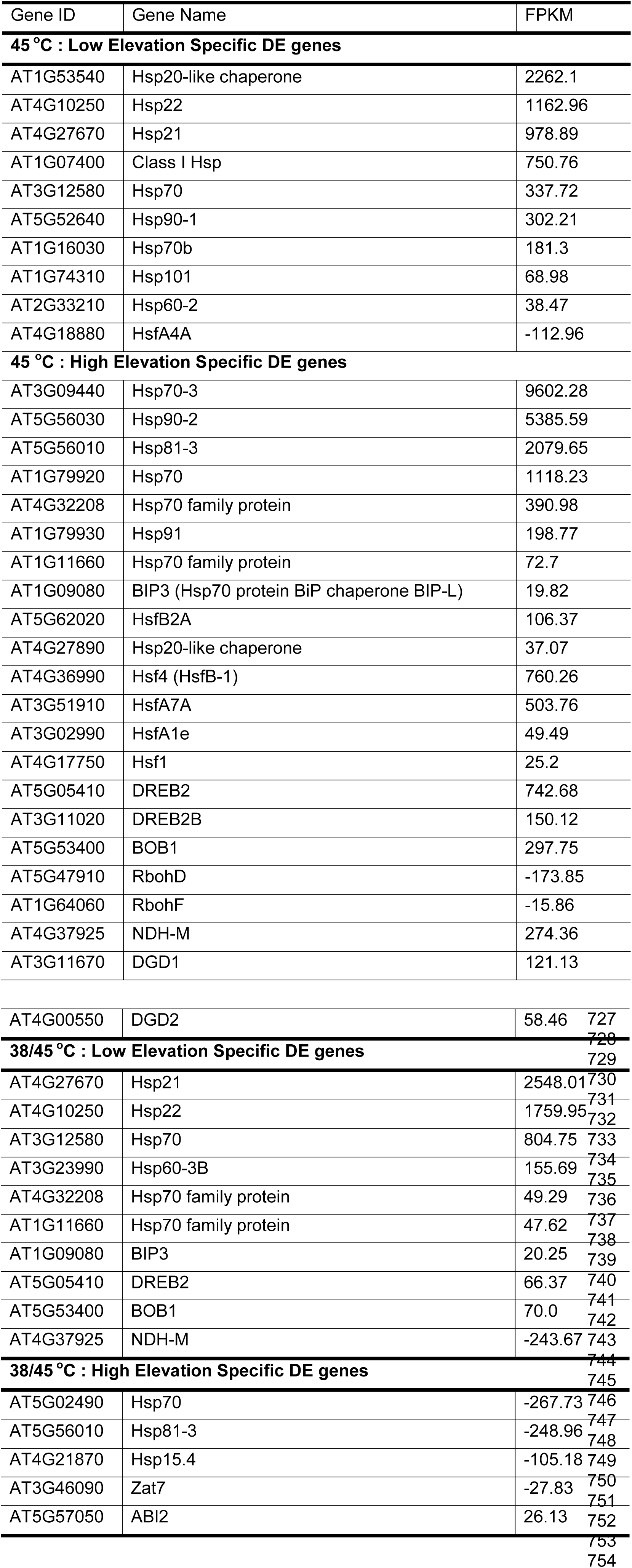
Currently known heat stress related elevation-specific DE genes in the two heat treatments

Note: FPKM, short for Fragments Per Kilobase Of Exon Per Million Fragments, is the normalized fold changes in gene expression when comparing high elevation plants to low elevation plants. Positive value means up-regulation, and negative value means down-regulation.

In 38/45°C, we found ten low elevation-specific and five high elevation-specific DE heat stress related genes. Low elevation plants showed up-regulation in small Hsps, Hsp60s and Hsp70s, and small Hsps showed much higher fold change than Hsp60s and Hsp70s. High elevation plants showed down-regulation in three Hsps (Hsp70, Hsp81-3 and Hsp15.4). Low elevation plants also showed up-regulation in DREB2 and BOB1 gene, increasing stress related genes. However, NDH-M gene was down-regulated in low elevation plants, potentially affecting photo-oxidation protection. In high elevation plants, Zat7 showed down-regulation. This might reduce the amount of antioxidant produced via the ROS pathway. High elevation plants also showed up-regulation in ABA signaling factor ABI2. In summary, in 38/45°C, low and high elevation plants were activated in both the Hsp/Hsf and the ROS pathway. Low elevation plants had up-regulation in all Hsps, especially small Hsps; high elevation plants had down-regulation in the Hsps. Low elevation and high elevation also adopted different genes in the ROS pathway.

### Acclimation-specific DE genes in low and high elevation plants

To uncover acclimation-specific DE genes in low and high elevation plants separately, we contrasted DE genes in 45°C with 38/45°C for plants from each elevation. The DE genes that were uniquely expressed in 38/45°C for plants from each elevation were acclimation-specific genes. We found 953 shared DE genes between the two heat treatments and 536 acclimation-specific DE genes for low elevation plants (Fig. 3a). Among the 536 acclimation-specific DE genes, 303 were up-regulated and 233 were down-regulated. We found 947 shared DE genes and 1025 acclimation-specific DE genes in high elevation plants (Fig. 3b). Among the 1025 acclimation-specific DE genes, 341 were up-regulated and 695 were down-regulated. High elevation plants showed more acclimation-specific DE genes than low elevation plants.

**Figure 3.**
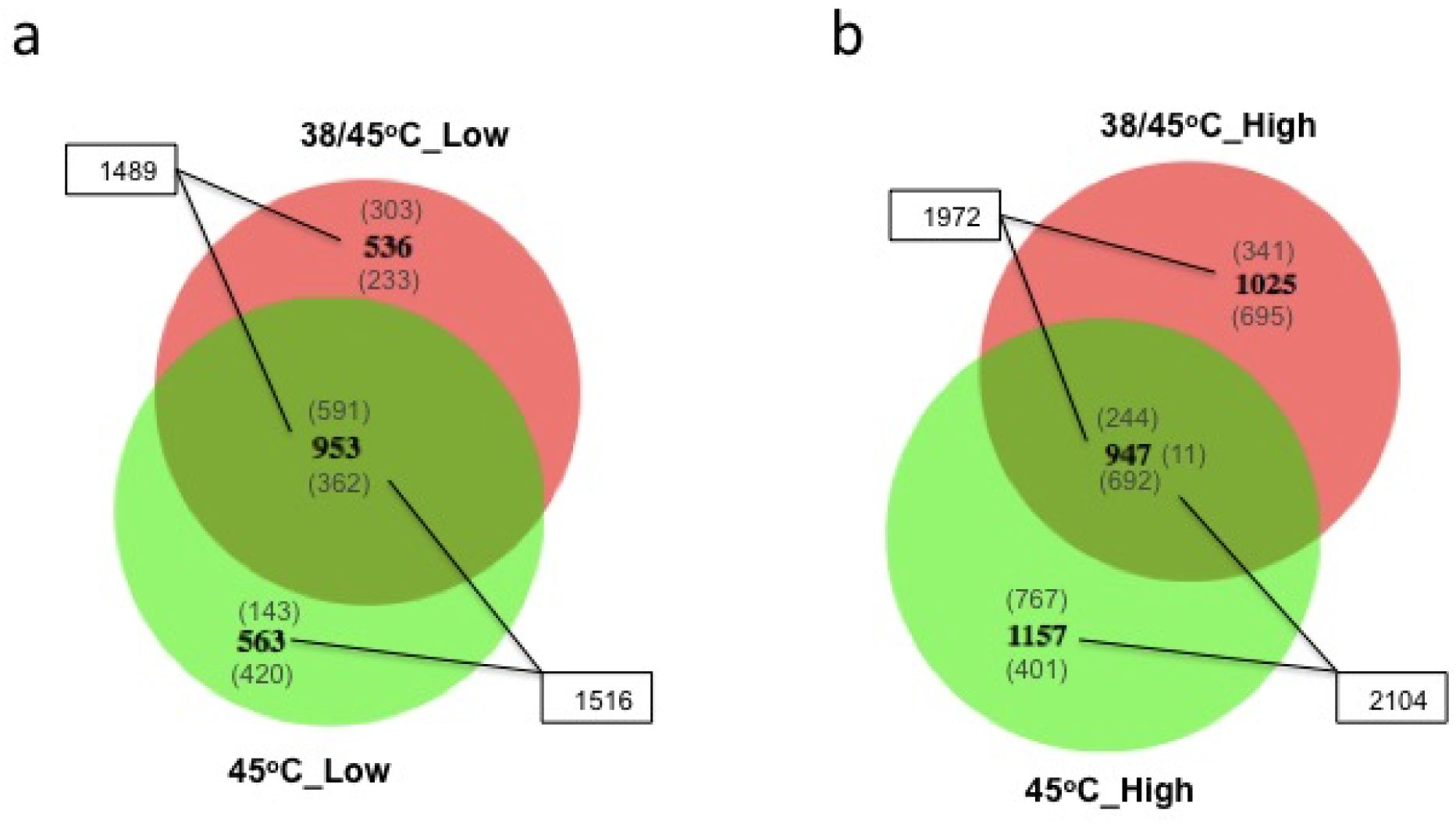
Venn diagrams comparing the number of DE (significantly differentially expressed) genes for the indicated treatment and elevation groups. In all cases DE genes are those differing significantly in expression compared to the same genes expressed in the corresponding 22°C control group. Bold numbers indicate total number of genes showing changed expression. Numbers in parentheses above the bold number indicate up-regulated genes and numbers below the bold indicate those that are down-regulated. The area of overlap of the two circles indicates the proportion of DE that are shared between the treatment and elevation groups. In the area of overlap, numbers in parentheses to the right of the bold numbers indicate DE genes that were shared but with directions of change between the compared groups, e.g. up-regulated in one group but down-regulated in the other group.

To further explore the acclimation-specific DE genes, we investigated the expression level of each gene in Table S1 from the above-mentioned acclimation-specific DE genes (Fig.2), for plants from each elevation (Table 3). Genes in Table S1 include genes involved in Hsp/Hsf pathway and ROS pathway. There were seven and eight acclimation-specific DE genes for low and high elevation plants respectively (Table 3). In the seven acclimation-specific DE genes in low elevation plants, only two Hsps, Hsp70 and BIP3, and one Hsf, HspA1e, were up-regulated. Two DREB genes also showed up-regulation. In the eight acclimation-specific DE genes in high elevation plants, six Hsps were up-regulated, with small Hsps having the largest magnitude of fold change. High elevation plants also experienced mostly down-regulation in the ROS pathway, such as Zat7. Hsps in high elevation plants also showed much higher magnitude of change than low elevation plants. The difference in expressed Hsps and other genes in ROS pathway showed that with acclimation, low elevation plants mainly adopted up-regulation in Hsp70s in Hsp/Hsf pathway and DREB genes in ROS pathway; high elevation plants adopted up-regulation in small Hsps, Hsp60s, Hsp70s, and Hsp101 in Hsp/Hsf pathway and ABA signaling in ROS pathway.

**Table 3.** Acclimation Specific heat stress related DE genes for low and high elevation plants

| Gene ID | Gene Name | FKPM |
| --- | --- | --- |
| **Acclimation specific stress related DE genes in low elevation plants** |  |  |
| AT1G11660 | Hsp70 family protein | 47.62 |
| AT1G09080 | BIP3 | 20.25 |
| AT3G02990 | HsfA1e | 18.38 |
| AT5G05410 | DREB2 | 66.37 |
| AT3G11020 | DREB2B | 46.27 |
| AT5G53400 | BOB1 | 70 |
| AT4G37925 | NDH-M | -243.67 |
| **Acclimation specific stress related DE genes in high elevation plants** |  |  |
| AT1G53540 | Hsp20-like protein | 2411.27 |
| AT1G07400 | Class I Hsp | 849.01 |
| AT1G16030 | Hsp70b | 249.9 |
| AT1G74310 | Hsp101 | 101.04 |
| AT2G33210 | Hsp60-2 | 72.38 |
| AT5G02490 | Hsp70 | -267.73 |
| AT3G46090 | Zat7 | -27.83 |
| AT5G57050 | ABI2 | 26.13 |

Note: FPKM, short for Fragments Per Kilobase Of Exon Per Million Fragments, is the normalized fold changes in gene expression when comparing high elevation plants to low elevation plants. Positive value means up-regulation, and negative value means down-regulation.

**Table 4.**
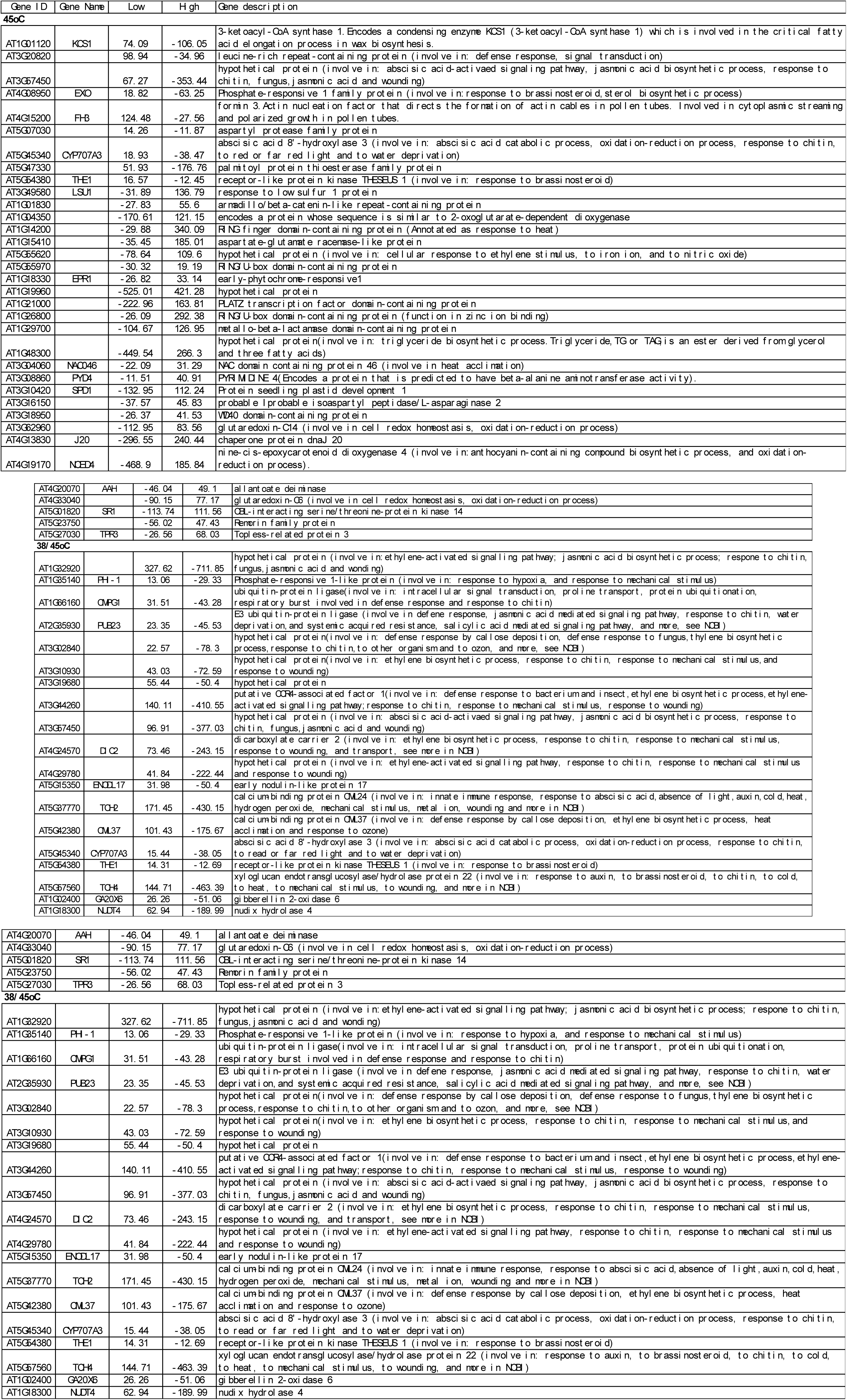
DE genes that were shared between low and high elevation plants but different directions.

### DE in both high and low elevation plants, but opposite directions of change

There were shared DE genes between low and high elevation plants that show opposite directions of change. There were 35 genes that were differentially expressed in the 45°C treatment (Fig.1a) and 19 in the 38/45°C treatment (Fig.1b) for which expression change was in opposite directions in the two elevation groups (Table 4). The functions of these genes mainly involve response to abiotic or biotic stress (such as heat, cold, chitin, ethylene stimulus, and wounding), signal transduction, biosynthetic processes, oxidation-reduction processes and cell redox homeostasis.

Of the 35 and 19 DE genes showing opposite directions of expression in the two elevation groups, three genes were included among both the 35 and 19 genes of this type from the two heat treatments: AT5G45340, AT5G54380, AT3G57450. Their functions primarily relate to abscisic acid(ABA)-activated signaling pathways, associated with response to abiotic or biotic stress. In the 35 shared DE genes at 45°C, nine DE genes showed up-regulation in the low elevation plants but down-regulation in the high elevation plants. The remaining 26 out of 35 DE genes showed down-regulation in the low elevation plants but up-regulation in the high elevation plants. However, all 19 shared DE genes at 38/45°C heat showed down-regulation in the low elevation plants but up-regulation in the high elevation plants. Among the 51 genes (that is: 35 + 19 – 3 overlapping), about 30 have been categorized as response to abiotic or biotic stresses (Table 4). These shared DE genes with opposite directions of change further showed the diversification in response to heat stress between low and high elevation populations.

## Discussion

When plants are exposed in high temperature, they not only experience heat stress, a secondary stress, oxidative stress, is also activated. Thus both genes in Hsp/Hsf pathway and in reactive oxygen species (ROS) pathway, including antioxidant and plant hormones, are produced (Qu *et al.* 2013). Here we compared the gene expression patterns for low and high elevation plants in NE Spain, in response to 45°C and 38/45°C treatments. High elevation plants had constitutively higher heat stress gene expression level, in both Hsp/Hsf and ROS pathway. In 45°C, only the Hsp/Hsf pathway was activated and small Hsps had the highest magnitude of change in low elevation plants; both Hsp/Hsf and ROS pathway were activated, with Hsp70 and Hsp90 showed the largest magnitude of fold change, in high elevation plants. in 38/45°C, low and high elevation plants were activated in both Hsp/Hsf and ROS pathway. Low elevation plants had up-regulation in all Hsps, especially small Hsps; high elevation plants had down-regulation in the Hsps. Low elevation and high elevation also adopted different genes in the ROS pathway. We also discussed shared genes between low and high elevation plants but with directions of change. This study indicates that low and high elevation plants have evolved adaptive divergence in heat stress response. The contrasting patterns of temperature variation in low and high elevation sites appears to have played a strong role in the evolution of divergent patterns of both pre-acclimation and direct exposure gene expression responses to high temperature stress.

### Population divergence in response to heat stress

Even when populations were not under heat stress, there was significant divergence in gene expression. This could potentially explain much of the phenotypic variation, such as flowering time, seed size, we documented previously in plants from the present study populations and others in this region (Montesinos-Navarro et al. 2012; Montesinos-Navarro et al. 2009; Montesinos-Navarro et al. 2011; Wolfe & Tonsor 2014). For example, low elevation plants flowers early but high elevation plants take a longer time to flower (Wolfe & Tonsor 2014). This can be explained by the expression of MADS box gene FLC and SOC1. FLC is a repressor of flowering (Michaels & Amasino 1999) and SOC1 promotes flowering (Lee & Lee 2010). FLC gene was up-regulated and the SOC1 was down-regulated in high elevation plants relative to low elevation plants, thus repressing flowering.

Natural populations of redband trout from desert sites showed the most uniquely differentially expressed transcripts and most abundant differentially expressed genes compared with populations from montane environment when exposed to severely high water temperatures (Narum & Campbell 2015). In response to a common thermal environment for intertidal snail *Chlorostoma funebralis*, more stress-responsive genes were observed in northern populations than southern populations (Gleason & Burton 2015). In the native environment of our samples, low elevation plants experience hot and dry climate while high elevation plants experience cold and wet conditions (Montesinos-Navarro *et al.* 2012; Montesinos-Navarro *et al.* 2009; Montesinos-Navarro *et al.* 2011; Wolfe & Tonsor 2014). Thus we hypothesize that low elevation plants potentially evolve to be more adapted to heat stress, through acclimation to more frequent high temperature events, but high elevation plants are more sensitive. Our data supports this hypothesis. High elevation plants expressed more elevation specific DE genes than low elevation plants in both heat treatments (Fig.1). In 45°C heat, high elevation plants showed more currently known heat stress related elevation specific DE genes than low elevation plants. However, with acclimation, low elevation plants showed up-regulation in Hsps but high elevation plants showed down-regulation in Hsps (Table 2). It is also worthwhile to notice that for high elevation plants in 38/45°C heat, there were 1088 elevation specific down-regulated DE genes, indicating substantial gene down-regulation involved (Fig. 1b).

Our phenotypic measures on the same set of biological replicates used in this study showed contrasting avoidance and tolerance strategies in a 45°C heat stress response. High elevation populations showed more avoidance, with lower rosette temperature at heat stress; and low elevation populations adopted more tolerance, i.e. a relatively higher photosynthetic rate (Zhang *et al.*, under review). Avoidance mechanisms include rosette angle and transpirational cooling, and tolerance mechanisms involve heat shock proteins, and plant hormones in the ROS (reactive oxygen species) pathway (Zhang et al under review). However, to the best of our knowledge, in our investigation on the current known heat stress related genes, the genes listed on Table S1 were all about mechanisms involved in tolerance, we did not find genes related with avoidance, in stress response. Low elevation plants had higher tolerance in response to 45°C heat, by expressing more DE genes (Fig. 1a, Table 2).

### Role of acclimation in heat stress response

Acclimation, from previous exposure to a sub-lethal high temperature, is an important adaptive mechanism and can enhance the ability to resist heat stress. Long-term acclimation can reach a new steady state with cost-effective strategies for stress response (Logan & Somero 2010).

When we compare 38/45°C with 45°C heat, we can look the genes specifically involved in acclimation. In 38/45°C, both low and high elevation plants showed fewer DE genes than the number observed in response to direct exposure to 45°C heat (Fig.2, Table 3). When we looked at the acclimation specific DE genes, low and high elevation plants had very different acclimation specific DE genes. Low elevation plants had 536 acclimation specific DE genes and high elevation plants had 1025 acclimation specific DE genes (Fig.2). For the currently known heat stress related DE genes among the acclimation specific DE genes, low elevation plants adopted two Hsps and one Hsf, but high elevation plants adopted six Hsps. Low elevation and high elevation plants also adopted very different genes in ROS pathways.

Hsp101 was previously reported to be the only Hsp that was necessary for heat tolerance (Hong & Vierling 2001). Here, we only found Hsp101 in low elevation plants in the 45°C heat, and in the high elevation plants in the 38/45°C heat. Hsp101 showed no up-regulation from 22°C - 34°C, but showed significant increase in 40°C compared with 34°C (Tonsor *et al.* 2008). When exposed to a 45°C heat stress, low elevation plants experienced average 38.2°C but high elevation plants experienced average 36.4°C because of avoidance mechanisms, such as transpirational cooling and inherent leaf angle change (Zhang *et al.* under review). The low temperature for high elevation plants, average 36.4°C, might not be high enough to activate Hsp101 expression. The difference in Hsp101 expression also showed very different mechanisms in stress response and acclimation between low and high elevation plants.

### Adaptive divergence of Hsps and Hsfs

We found that populations from low elevation, a hot and dry environment compared to the environment of high elevation plants (Montesinos-Navarro *et al.* 2009; Montesinos-Navarro *et al.* 2011; Wolfe & Tonsor 2014), showed lower Hsp101 expression than populations from high elevation, cold and wet environment in the 45°C heat treatment, although the difference did not reach statistical significant because of high variation in expresion at high temperature (Zhang *et al.* 2015a). This is in accordance with our gene expression data here. We found that low elevation plants showed fewer DE genes for Hsps in the heat treatments, compared to high elevation plants.

A similar pattern has been seen in redband trout (Narum *et al.* 2013), common killifish (Fangue *et al.* 2006) and intertidal snail (Gleason & Burton 2015). Studies on redband trout also showed lower Hsp expression observed in desert strains compared to montane strains (Narum & Campbell 2015; Narum *et al.* 2013). Studies on common killifish showed significantly greater Hsp70-2 gene expression in the northern than the levels observed in southern killifish populations (Fangue *et al.* 2006). Hsp70 is involved in negative regulation of heat stress response (Morimoto 1998). These studies combined indicate populations from warm environments might have evolved heat tolerance mechanisms with lower costs than Hsps production. However, studies in common killifish also showed other Hsps, such as Hsp70-1, or Hsp90, had different patterns in response to heat stress, suggesting that Hsps have complex networking patterns in heat stress responses. In response to a common thermal environment for intertidal snail *Chlorostoma funebralis*, the two regions also showed important differences in the genes that were up-regulated. Hsp70s were significantly increased in the northern populations while Hsp40s were significantly up-regulated in the southern populations (Gleason & Burton 2015). This is also in accordance with our findings on Hsps in this study, in which we saw various magnitudes of gene expression and different Hsps in low and high elevation plants (Table 2).

### Adaptive divergence in natural populations: ROS pathway

Although Hsp/Hsf pathway is still the major differentiated pathway between low and high elevation plants in heat stress response, other genes involved in ROS pathway also played a significant role in each elevation plant. Previous heat stress studies showed DE genes involve Hsps, and genes involved in ubiquitination and proteolysis (Schoville *et al.* 2012), as well as genes involved in oxygen transport, protein synthesis, folding and degradation in Saccharina japonica and catfish (Liu *et al.* 2013a; Liu *et al.* 2013b). Abscisic acid, salicylic acid, hydrogen peroxide and ethylene related signaling pathways are also involved in heat stress response (Larkindale *et al.* 2005; Larkindale & Huang 2005).

In elevation specific DE genes and acclimation specific DE genes, we found DE genes involved in ROS pathway, such as DREB2 and ABI gene (Table 2, Table 3). The shared DE genes with directions of change in gene expression in low vs. high elevation plants showed genes involved in ubiquitination (e.g., AT1G14200, AT5G55620), abscisic acid catabolic process (e.g., AT5G45340) and ethylene biosynthetic process or ethylene activated signaling pathway (e.g., AT4G24570, AT4G29780) (Table 4). Heat stress response is a complex process and it will certainly need more effort to clarify the genes involved and how they interact to determine phenotypic responses.

In nature, plants often face a combination of several stresses. However, plants’ response to stress combinations cannot be directly predicted from the response in each single stress (Rizhsky *et al.* 2002; Rizhsky *et al.* 2004b). The next steps of research should focus on two areas. One is to understand the cross-talk among various stress responses; the second is to understand the evolution of heat stress response and acclimation in plants from various climates. Since plants originate from different climates, they experience very different patterns of stress combination, thus they evolve differently in stress response. Looking into the agriculturally important stress combinations from the stress matrix (Mittler 2006) is the next challenge.

## Sources of funding

Funding was provided by US National Science Foundation Grant IOS-1120383 to SJT and xx to EV.

## Contributions by authors

NZ, EV and SJT designed the experiment. NZ performed the experimental work. NZ performed analyses. All co-authors discussed and interpreted results. Writing was done by NZ, EV and SJT.

## Acknowledgement

We are grateful to Tonsor Lab manager Tim Park for excellent technical assistance. SJT is extremely grateful to F. Xavier Picó of Estación Biológica de Doñana, Seville, Spain for introduction to the Spanish *Arabidopsis* system and for many hours of friendship during field characterization and collection of the populations.

## Supplementary Materials

**Table S1.** Alignment summary of RNA-seq data for the 24 samples to *Arabidopsis thaliana* (Tair 10) transcriptome with Tophat2.

| Sample | Elevation group | Treatment | Number Raw Reads | Number Mapped Reads | % of total mapped |
| --- | --- | --- | --- | --- | --- |
| HOR7 | Low | 22°C | 27736919 | 24308845 | 87.6 |
| HOR7 | Low | 45°C | 27735098 | 24823054 | 89.5 |
| HOR7 | Low | 38-45°C | 34285786 | 26806810 | 78.2 |
| PIN9 | Low | 22°C | 35250112 | 31290257 | 88.8 |
| PIN9 | Low | 45°C | 27414384 | 24493775 | 89.3 |
| PIN9 | Low | 38-45°C | 22861792 | 8279162 | 80.0 |
| RAB4 | Low | 22°C | 24956107 | 22166537 | 88.8 |
| RAB4 | Low | 45°C | 24910779 | 22711036 | 91.2 |
| RAB4 | Low | 38-45°C | 27280627 | 21763377 | 79.8 |
| SPE6 | Low | 22°C | 26441088 | 23290731 | 88.1 |
| SPE6 | Low | 45°C | 27479845 | 24828659 | 90.4 |
| SPE6 | Low | 38-45°C | 35813970 | 28071470 | 78.4 |
| BIS8 | High | 22°C | 24102301 | 21236376 | 88.1 |
| BIS8 | High | 45°C | 27300909 | 24737693 | 90.6 |
| BIS8 | High | 38-45°C | 31933204 | 25013914 | 78.3 |
| PAL6 | High | 22°C | 26127566 | 23085888 | 88.4 |
| PAL6 | High | 45°C | 25747254 | 23168349 | 90.0 |
| PAL6 | High | 38-45°C | 32030973 | 25127204 | 78.4 |
| PAN8 | High | 22°C | 26737620 | 23502729 | 87.9 |
| PAN8 | High | 45°C | 26428191 | 23708370 | 89.7 |
| PAN8 | High | 38-45°C | 30941863 | 24415423 | 78.9 |
| VIE4 | High | 22°C | 30350162 | 26693836 | 88.0 |
| VIE4 | High | 45°C | 26457362 | 23935965 | 90.5 |
| VIE4 | High | 38-45°C | 27106330 | 21442774 | 79.1 |

Note: The summary was based on alignment results after trimming out the first and last 15bp for each 100bp reads.

**Table S2.** Currently known heat stress related genes investigated in this study.

| Gene family | Gene description |
| --- | --- |
| **Hsp-Hsf pathway:** |  |
| Hsfs | Heat shock factors |
| Hsps | Heat shock proteins |
| **ROS pathway:** |  |
| Zat family<br>(Zat7, Zat10, Zat12) | Zinc transporter family protein, responds to diversified stress, including heat stress. |
| WRKY family<br>(WRKY25) | WRKY transcription factor, involve in response to heat and other stress. |
| AP2/ERF family<br>(DREB2A, DREB2B, ETR1, EIN2) | Plant specific transcription factor, activates the expression of abiotic stress-responsive genes. |
| MBF1c | Highly conserved transcriptional co-activator, involve in thermotolerance. |
| RBoh | ROS (reactive oxygen species) signal amplifier. |
| CBK3 | Important component of Ca<sup>2+</sup> -regulated heat stress signal transduction pathway, downstream of CaM, which regulates Hsps expression. |
| BOB1 (BOBBER1) | A small Hsp with a thermotolerance role at high temperature. |
| ABA signaling (ABI1, ABI2) | Reduced survival after heat stress in these mutants, however the accumulation of Hsps was not affected. |
| NDH1 | High heat-inducible and provide protection against photo-oxidation. |
| PP7 | Encodes a nuclear localized serine/threonine phosphatase that appears to be regulated by redox activity and is a positive regulator of cryptochrome mediated blue light signalling. |
| BI1 | BAX inhibitor 1, Functions as an attenuator of biotic and abiotic types of cell death. |
| UVH6 | A negative regulator of the common stress response induced by UV damage and heat. |
| VPS53 | Involved in vesicle trafficking, heat stress sensitive gene. |
| CTL1 | A chitinase-like protein, required for acquired thermotolerance, salt stress and development. |
| FtsH11 | Chloroplastic FtsH11 protease, associated with strongly reduced photosynthetic capacity after heat stress. |
| DGD1 | Digalactosyldiacylglycerol synthase 1, associated with strongly reduced photosynthetic capacity after heat stress. |
| TU8/TFL2 | Terminal Flower 2, which shows Hsp90 reduction in the tu8 mutant. |

Note: the above gene list is summarized from three review papers (Kotak et al. 2007, Ahuja et al. 2010, Qu et al. 2013). Gene functions were further confirmed from the NCBI website.

